# Developmental Patterning of Irritability Enhances Prediction of Psychopathology in Pre-adolescence: Improving RDoC with Developmental Science

**DOI:** 10.1101/2020.04.30.070714

**Authors:** Katherine S. F. Damme, Lauren S. Wakschlag, Margaret J. Briggs-Gowan, Elizabeth S. Norton, Vijay A. Mittal

## Abstract

Research has demonstrated the transdiagnostic importance of irritability in psychopathology pathways but the contribution of developmentally-unfolding patterns has only recently been explored. To address this question, irritability patterns of 110 youth from a large and diverse early childhood cohort were assessed at preschool age and at school age (∼2.5 years later) with a dimensional irritability scale designed to capture the normal:abnormal spectrum. Participants then returned at Pre-adolescence (∼6 years later) for an assessment with a structured clinical interview (internalizing/externalizing symptoms) and a magnetic resonance imaging scan. When only preschool age irritability was considered, this was a transdiagnostic predictor of internalizing and externalizing symptoms. However, a model including both preschool and school age irritability provided a more nuanced picture. A high preschool and decreasing school age profile of irritability predicted elevated pre-adolescence *internalizing* symptoms, potentially reflecting emerging coping/internalizing behavior in pre-adolescence. In contrast, a stable irritability profile across these timepoints predicted increased pre-adolescence *externalizing* symptoms. Further, preschool irritability (a period of rapid growth) did not predict pre-adolescent gray matter volume abnormality, an indicator of transdiagnostic clinical risk. However, irritability at school age (when gray matter volume growth is largely finished) demonstrated an interactive effect among regions; increased school age irritability predicted reduced volume in pre-adolescence emotional regions (e.g., amygdala, medial orbitofrontal cortex) and increased volume in other regions (e.g., cerebellum). Expanding the impact of RDoC’s approach yielding transdiagnostic phenotypes and multiple units of analysis, a developmentally informed approach provides critical new insights into the complex unfolding of mechanisms underlying emerging psychopathology.

## Introduction

The National Institute of Mental Health’s Research Domain Criteria (RDoC) initiative (Cuthbert & Insel, 2013) provides a framework for studying psychopathology by defining several primary (and a number of facet and sub-facet) transdiagnostic areas of dysfunction, as well as providing metrics of units, ranging from cells to complex behaviors (with brain circuits placed centrally as a target of many level of analyses) that make up these foundational elements of psychopathology (Cuthbert & Insel, 2013). What is missing from the current RDoC framework is the incorporation of core methods and concepts from the vibrant field of developmental science. When applied in a developmental framework, RDoC can be even more effective at clarifying mechanisms and informing treatment (Casey et al., 2014; Mittal & Wakschlag, 2017). A developmental approach may have significant potential for understanding the predictive potential of one of the earliest measurable behavioral signs of psychopathology: irritability.

Emerging psychopathology is evident early in life (Angold & Egger, 2007; Carter et al., 2013; Mittal & Wakschlag, 2017; Wakschlag et al., 2010). This early clinical phenomenology has been defined by atypicality from normative development, while considering neurodevelopmental patterns within developmental context. This approach has elucidated unfolding patterns of symptoms that predict later psychopathology and provide insight into etiology and inform developmentally-based interventions (Cicchetti & Sroufe, 2000; Mittal & Wakschlag, 2017; Wakschlag et al., 2010). Within this context, a wealth of evidence demonstrates that atypical patterns of early irritability are predictive of emerging and subsequent psychopathology (Brotman et al., 2017; Carter et al., 2013; Fishburn et al., 2019; Hawes et al., 2019; Humphreys et al., 2019; Kessel et al., 2020; Smith et al., 2019; Wakschlag et al., 2010, 2015a, 2019). Despite the promise of irritability to predict psychopathology, it is not yet clear how a general behavioral phenotype of excessive irritability translates into diverse, specific symptoms of psychopathology. This question likely remains because our current understanding of the dynamics of emerging psychopathology currently relies on snapshots of individuals at a single time points to predict outcomes, creating a noisy picture of these dynamic processes (Kaczkurkin, Moore, et al., 2019; Kjeldsen et al., 2014; Mittal & Wakschlag, 2017; Wakschlag et al., 2015a). Although most prior work examines irritability as a snapshot, our prior work indicates that accounting for longitudinal variation improves clinical prediction (Wakschlag et al., 2015a). To our knowledge, however, no studies have examined the impact of accounting for developmental patterning of irritability in prediction of neural outcomes.

Although some previous work has found important links with early irritability, psychopathology and aberrant, regional gray matter volumes (Andre et al., 2019; Goodkind et al., 2015; Kaczkurkin, Moore, et al., 2019; Kaczkurkin, Park, et al., 2019; Leibenluft, 2017; Pagliaccio et al., 2018), developmentally-driven studies that incorporate patterns of irritability with tools such as neuroimaging have been scant. The current study examines irritability at two critical developmental periods, preschool age and early school age (Manning et al., 2019; Wakschlag et al., 2017) to examine patterns predictive of pre-adolescent psychopathology and gray matter volume abnormalities.

Irritability is characterized as a dispositional tendency for an individual to respond to blocked goal attainment with anger (Wakschlag et al., 2017). Although some level of irritability is normative, atypical irritability patterns reflect intensity and dysregulation that is distinct from the expected ability in developmental context (Wakschlag et al., 2015a). Atypical irritability patterns robustly predict internalizing/externalizing psychopathology across the lifespan (Leibenluft, 2017; Stringaris, 2011; Wakschlag et al., 2019). Irritability thus may also present an opportunity to understand how general risk for psychopathology may interact with a confluence of developmental factors to manifest as specific symptoms of psychopathology in adolescence and adulthood (Kaat et al., 2019; Kjeldsen et al., 2014; Mittal & Wakschlag, 2017; Smith et al., 2019; Wakschlag et al., 2010, 2014). Advances in developmental measurement science now enable characterization of its normal:abnormal spectrum within developmental context. Irritability at early school age, for example, may be impacted by neurodevelopmental contexts such as the co-occurring maturation of critical social, emotional, and regulatory gray matter volumes (Kaczkurkin, Moore, et al., 2019). These influences on irritability may identify unique targets for early intervention and prevention of the development of psychopathology.

During early development self-regulatory capacities are evolving and have been shown to be predictive of later psychopathology (Wakschlag et al., 2010, 2017). Critically, this is also a time of development of basic self-control, evident in verbal negotiation, delay of gratification, frustration tolerance, and behavioral flexibility (Wakschlag et al., 2005), which may reflect underlying neurodevelopment. The preschool period is also one of shifting environmental and social expectations, in which parents have increasing expectations and peers begin to play a larger role in socialization (Wakschlag et al., 2005). This period also marks a shift in neurodevelopment, as cell and synaptic division peak; increased synaptic pruning, arborization, and myelination alter gray matter volumes, which peak in pre-adolescence (Giedd et al., 1999, 2012; Kaczkurkin, Moore, et al., 2019). Despite this complex developmental context, preschool irritability can be measured reliably and is predictive of future psychopathology (Hawes et al., 2019; Kessel et al., 2020; Smith et al., 2019; Wiggins et al., 2020; Wiggins et al., 2018).

In addition to preschool, early school age is a critical period for development of emotional regulation (Briggs-Gowan et al., 2006). Changes in irritability from preschool may reflect emotion development and regulation strategies. These developments presage internalizing and externalizing symptoms in pre-adolescence, when psychopathology is emerging and stabilizing (Class et al., 2019; Hawes et al., 2019; Jirsaraie et al., 2019). Additionally, early school age is a time of rapid gray matter volume maturation. These changes in gray matter volume may provide critical insight into development of psychopathology, as volumes also peak in pre-adolescence (Giedd et al., 1999, 2012), which is when symptoms stabilize. In adulthood, gray matter volume is predictive of transdiagnostic psychopathology, i.e., both internalizing and externalizing symptoms (Aoki et al., 2014; Davis et al., 2018; Haijma et al., 2013; Shang et al., 2014; Wise et al., 2017). Critically, changes in gray matter volume from preschool to a peak in pre-adolescence may reflect critical neurodevelopmental trajectories and provide insight into how irritability may reflect and impact neurodevelopment (Kaczkurkin, Moore, et al., 2019). Irritability during this time may reflect the underlying neurodevelopment of emotional, social, and cognitive (i.e., executive control) systems that would be apparent in gray matter volumes at pre-adolescence, when specific externalizing and internalizing disorders are evident (Tseng et al., 2018).

Irritability indicates future psychopathology, but it is unknown how variations over development, potentially reflecting multiple unique pathways to common, modifiable, and broad symptoms of psychopathology (i.e., internalizing and externalizing) or resulting biological markers of general risk for psychopathology (i.e., gray matter volume). To investigate these questions, the current study examines irritability at multiple time points (preschool age, school age) to predict clinical outcomes (internalizing and externalizing symptoms) and gray matter volume in pre-adolescence. In a second model, the current study examines whether the addition of a second time point in early school age adds insight into prediction internalizing and externalizing symptoms at pre-adolescence. Additionally, the current study will examine if preschool and school age irritability, when rapid neuromaturation of gray matter volume occurs, will relate to differences in regional gray matter volumes at pre-adolescence.

## Methods and Materials

### Participants

The current study uses a subsample of data from a larger longitudinal dataset, the Multidimensional Assessment of Preschoolers Study (MAPS). The MAPS study is a longitudinal study enriched for psychopathology via oversampling for disruptive behavior and violence exposure (for details see Briggs-Gowan et al., 2019; Wakschlag et al., 2015a). Within the larger MAPS study, 425 preschoolers at baseline participated in an intensive laboratory-based study; of these participants 355 (83.5%) were seen again at early school age, and 303 (71.3%) returned for a lab visit at pre-adolescence. Of the 303 participants who attended lab visits at the pre-adolescence wave, 197 were invited to participate in the MRI; 13 of those pre-adolescence subjects were ineligible due to MR contraindications (e.g. braces); out of the remaining 184, there were 129 consented and 120 participated. As a result, the current neuroimaging subsample of 120 youth were selected for participation in this neuroimaging sub-sample using a sample frame designed to yield a sample with variation in early irritability and low concern for others along the full spectrum (low to high), based on mother-reports on the dimensional measure the Multidimensional Assessment of Profile of Disruptive Behavior (MAPDB) at the preschool wave. Prior to the baseline at the preschool age, participants were assigned sampling scores indicating their priority for invitation into the subsample. Those who did not participate in PA or declined MRI were replaced by the next participant according to rank order. Consistent with the sampling strategy, 41% of the MRI sample had high levels of irritability at preschool (above the 80th percentile in a normative sample). Ten individuals were excluded from the analyses due to motion artifacts in the data resulting in a final sample size of 110 (44% female). Although the project focuses on continuous outcomes (an RDoC priority), it is important to recognize that many of the late childhood cases are expressing symptoms at a clinically relevant level, and in some cases, these youth already meet criteria for an internalizing (8.65% that meet diagnostic criteria for depression, separation anxiety, and/or generalized anxiety) or externalizing (21.15% that meet diagnostic criteria for attention deficit hyperactivity disorder, conduct disorder, and/or oppositional defiant disorder) disorders. Critically, this sub-sample with an MRI was not significantly different from the larger sample previously reported in the literature in terms of sex (*p*=.11), race (*p*=.43), income (*p*=.85) age (*p*=.09), or irritability at preschool age (*p*=.58). This subsample included irritability scores at Preschool Age (age *M*=4.17, *SD*=.77) and School Age (*M*=6.95, *SD*=.99), as well as MRI and structured clinical interview at pre-adolescence (*M*=9.23, *SD*=.43). The sample was ethnically and racially diverse: 51.0% African American, 28.8% Hispanic and 20.2% Caucasian.

### Clinical Assessments

#### Irritability

Irritability was assessed at both preschool and school age with the Temper Loss scale of the Multidimensional Assessment Profile of Disruptive Behavior (MAP-DB; Kaat et al., 2019; Wakschlag et al., 2014). The MAP-DB was specifically designed to characterize the normal:abnormal spectrum of irritability within developmental context. This includes 22 items that account for the variation in intensity, context, and dysregulation, as well as objective frequency ratings (e.g. times per week), which provide developmentally meaningful information about rates of occurrence to reduce error inherent in subjective judgements, e.g., often, rarely (Wakschlag et al., 2017). IRT methods were used to generate irritability severity along four dimensions (temper loss, low concern, aggression, non-compliance). The resulting irritability scores indicate the relative position of the response in a four dimensional distribution of responses whereby a higher number reflects an increase in severity and the endorsement of items not frequently endorsed, and a lower score indicates that the irritability levels were lower than the irritability in the independent reference sample (for more informaiton see Wakschlag et al., 2014).

#### Clinical Assessment of Internalizing and Externalizing Symptoms at Pre-adolescence

At the EA time point, participants were assessed via symptom counts via parent interview with the Kiddie Schedule for Affective Disorders and Schizophrenia–Present and Lifetime Version (K-SADS-PL; Kaufman et al., 2000). Items on the K-SADS-PL items are scored from 0 to 3-point where a score of 0 indicate the symptom is not present, 2 is subthreshold levels of symptomatology, and 3 indicates reaching clinical a clinically relevant symptom on that item. For the current analyses, total number of symptoms were counted for externalizing and internalizing disorders.

#### MRI Acquisition and Processing

MRI data were collected on a 3-Tesla Siemens Tim Trio scanner (Erlangen, Germany) using a standard 32-channel head coil. Structural images were collected with an echo planar imaging navigated T1-weighted 3D magnetization prepared rapid gradient multi-echo (MPRAGE) sequence (sagittal plane; repetition time (TR)=270ms; echo times (TE)=1.69, 3.5, and 5.41; GRAPPA parallel imaging factor 2; 1 mm^3^ isotropic voxels, 192 interleaved slices; FOV=256mm; flip angle=7°). MRI data were visually inspected for quality of gray matter segmentation and motion artifacts (Iscan et al., 2015). Freesurfer v.6.0,(Fischl, 2012) software automatically labeled anatomical segmentations according to the Desikan atlas (Desikan et al., 2006) to calculate gray matter volumes in region of interest across the whole brain.

## Results

### Participants

Irritability scores ranged from −1.64 to 3.03 (*M*=.28, *SD*=.11) at the preschool age time point and from −1.64 to 2.28 (*M*=−.12, *SD*=.09) at the school age time point. From preschool to school age, 52.7% of the sample showed an increase or stable irritability score and 47.2% of the sample showed a decrease in irritability, *Figure 1A*. The KSADs internalizing symptoms scores ranged from 0-16 (*M*=1.83, *SD*=3.14) and externalizing symptoms scores ranged from 0-24 (*M*=3.91, *SD*=5.40).

**Figure 1.**
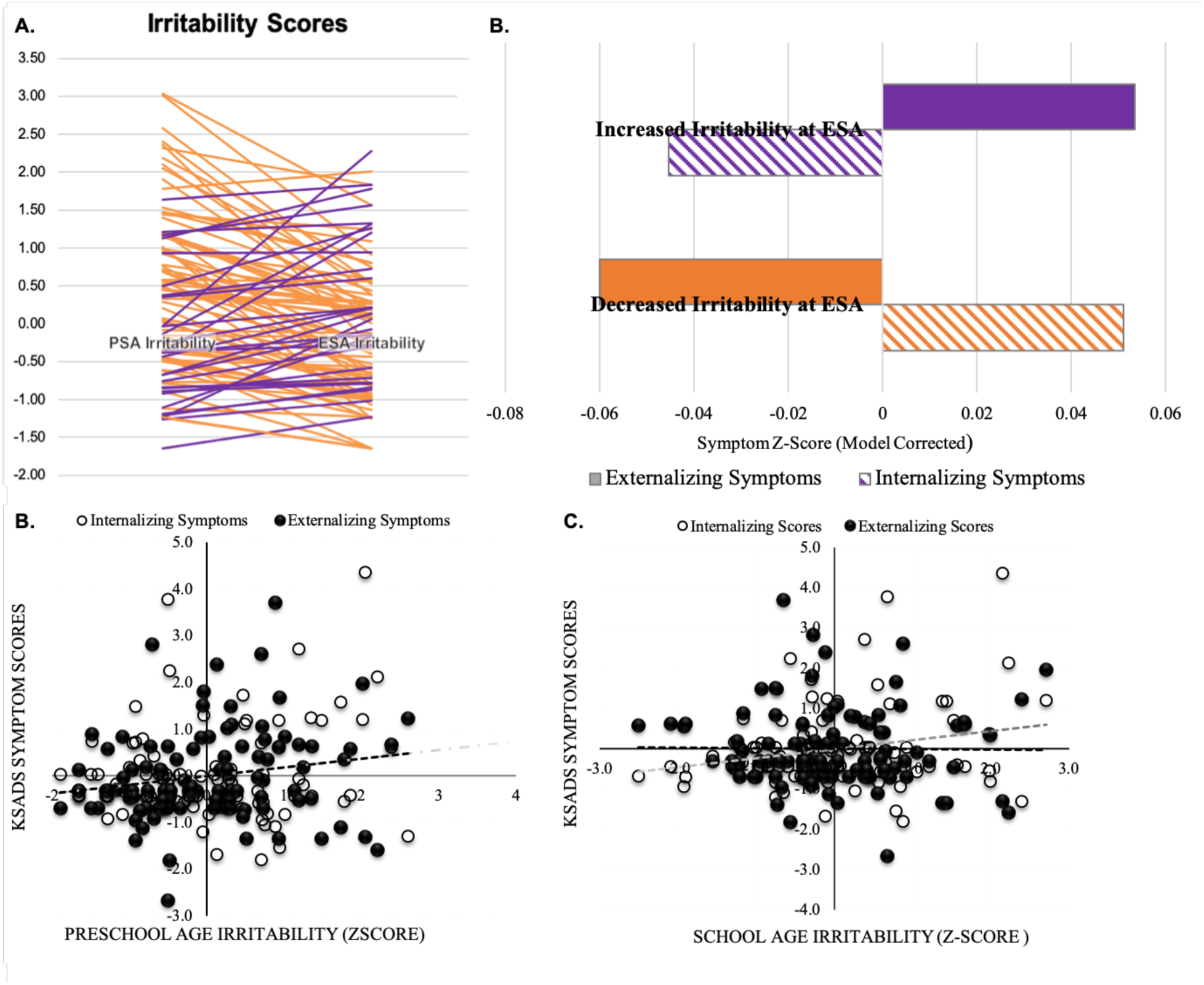
Early Irritability Scores Related to Pre-adolescent Symptoms of Psychopathology: A. Preschool Age Irritability Showed a Positive Relationship to Internalizing and Externalizing Symptoms at Pre-adolescent, B. Irritability Trajectories (From Preschool to School Age) Related to Distinct Pre-Adolescent Symptom Preschool Age, C. Irritability Related to School Age Internalizing and Externalizing symptoms, D. School Age Irritability Related to School Age Internalizing and Externalizing symptoms

### Preschool Irritability Alone Predicting Pre-Adolescent Symptoms

A general linear model defining the dependent variable as irritability at the preschool age time point alone was related to the internalizing and externalizing counts of total number of symptoms at pre-adolescence accounting for child sex. The omnibus test of model was significant, *F*(1,105)=6.57, *p*<.001 *partial-eta*^*2*^=.163. Irritability scores at the preschool are time point related to internalizing symptoms scores at the pre-adolescence time point, *F*(1,105)=4.38, *p*=.041, partial eta^*2*^=.04. Irritability scores at the preschool time point related to externalizing symptoms scores at the pre-adolescence time point, *F*(1,105)=5.01, *p*=.026, partial-eta^*2*^=.05, *Figure 1c*.

### Irritability at Multiple Early Time Points Predicting Pre-Adolescent Symptoms

In a repeated-measures general model, preschool and school age irritability scores were defined as within-subject measures, predicting externalizing and internalizing sum scores at pre-adolescence, accounting for sex. There was a main effect of irritability predicting internalizing scores at pre-adolescence, *F*(1,105)=8.38, *p*=.005, partial-eta^2^=.08, and this effect did not vary by time point, *p=*.30. Post-hoc analyses revealed that high irritability (regardless of time point) predicted increased pre-adolescent externalizing symptoms (partial-*r’*s =.35-.38). There was a significant time point by internalizing symptoms interaction, *F*(1,105)=4.52, *p*=.03, partial-eta^*2*^=.05, *Figure 1d*, such that high irritability at preschool age (partial-*r*=.21), and decreased (reduced when accounting for preschool age) irritably at school age (partial-*r*=−.13) predicted elevated pre-adolescent internalizing symptoms, *Figure 1e*, but there was no main effect of irritability across time points predicting internalizing symptoms at pre-adolescence, *p*=.18.

### Irritability at Multiple Early Time Points Predicting Pre-Adolescent Cortical Grey Matter Volumes

In a repeated-measures general linear model, 43 brain region volumes (9 subcortical regions and 34 cortical regions) nested in two hemispheres (left and right) as within-subject variables were related to the preschool and school age irritability ratings as between-subject variables, accounting for variance in volume related to sex and age at scan. There were no main effects of preschool age irritability scores (*p*=.45) or interactive effects (*p’*s>.31) on gray matter volumes. In contrast, irritability at pre-adolescence showed an interactive effect of school age irritability by regions, *F*(42,4158)=1.88, *p*=.001, but no main effect on gray matter volume (*p*=.39). This regional specificity demonstrated that increased school age irritability predicted reduced volume (partial-*r’*s=−.19 to −.13) in pre-adolescent emotional regions (e.g., amygdala, medial orbitofrontal cortex, hippocampi) and increased volume (partial-*r’*s=.19 to .13) in other regions (e.g., cerebellum), *Figure 2*.

**Figure 2.**
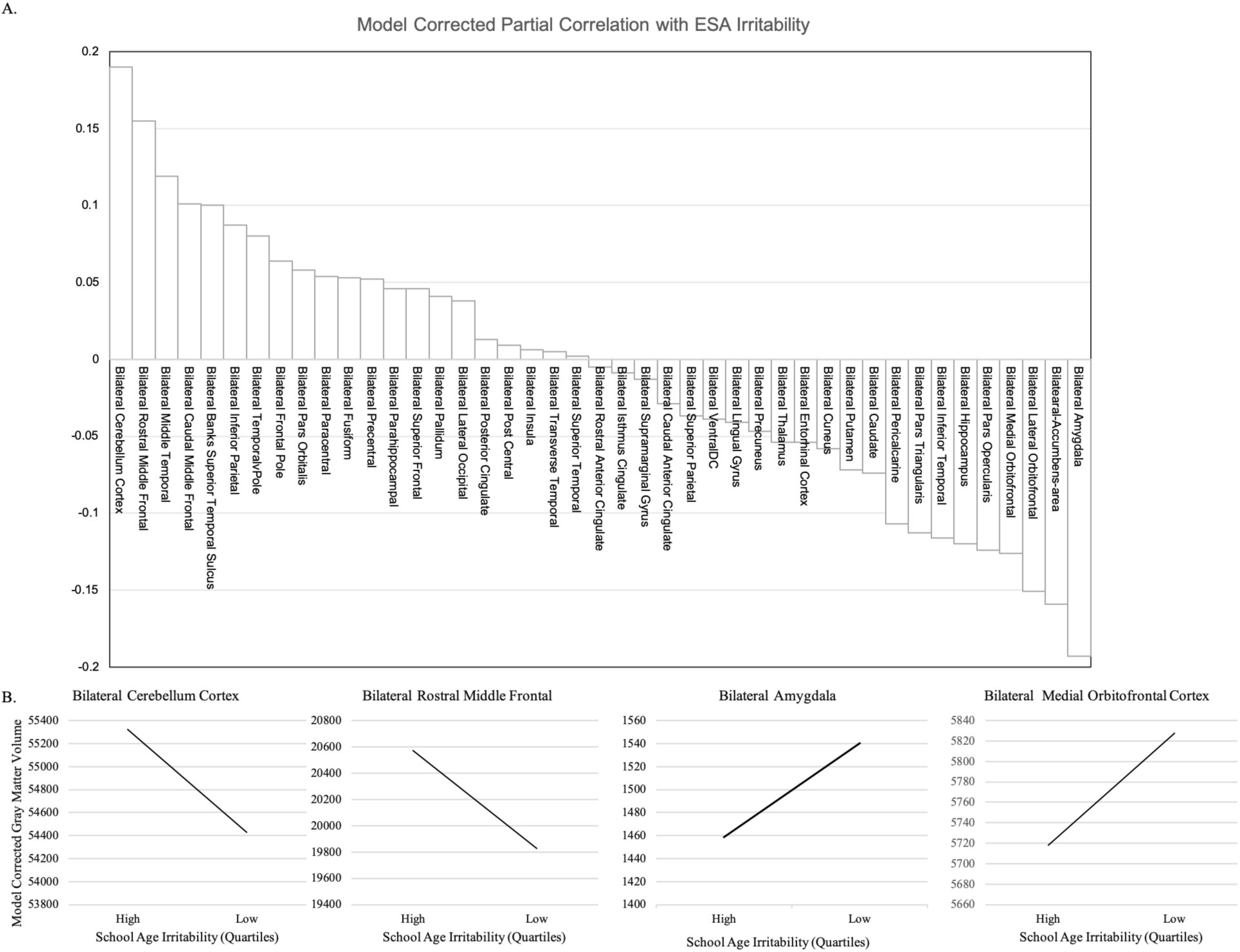
Early School Age Irritability Scores Related to Early Adolescent Cortical Volumes: A. Depicts the Model Corrected Partial Correlations between Cortical Volume and School Age Irritability Scores B. Depicts Highest Effect Sizes Model Corrected Volumes Related to School Age Irritability

## Discussion

The current findings elucidate pathways by which irritability develops into a multi-finality of adolescent clinical and neural outcomes (Kaczkurkin, Moore, et al., 2019; Kjeldsen et al., 2014; Mittal & Wakschlag, 2017; Pagliaccio et al., 2018). First, the preschool measurement provided critically important insight into general risk for psychopathology, indicating that early irritability predicted both internalizing and externalizing symptoms in pre-adolescence. Adding the early school age time point provided critical specificity to these findings. Irritability at school age predicted specific relationship to internalizing; elevated irritability at both time points related to higher externalizing symptoms in pre-adolescence, whereas reductions in school related to internalizing at pre-adolescence. Critically, these findings emphasize the importance of examining the developmental patterning of irritability for improving specificity in prediction (Mittal & Wakschlag, 2017; Wakschlag et al., 2012). As we only incorporated the two childhood timepoints available in the MAPS study, we do not know whether these findings are anchored in the specific developmental periods studied (from preschool to school age) or reflect the need to account for change over time regardless of developmental period. In terms of brain measures, school age, but not preschool age, irritability predicted variability in gray matter volumes at pre-adolescence, potentially reflecting underlying neuromaturational processes co-occurring/relating to changes in irritability. In summary, the relationship of irritability to clinical and neural outcomes is dynamic over development and varies in its predictive utility. Elevated irritability is broadly predictive early, but it is the developmental profile of irritability patterns that predicts the multi-finality of specific symptoms and underlying neurodevelopment in school age (Pagliaccio et al., 2018; Wakschlag et al., 2015a).

Elevated irritability at preschool is a strong predictor of a general psychopathology in pre-adolescence, which emphasizes the utility of this developmental time point as a clear marker of trait tendencies (Wakschlag et al., 2015b). The pattern of irritability over development, however, provides critical insight, unmasking a latent specificity of risk for psychopathology in irritability during early school age (Briggs-Gowan et al., 2016). This second finding illuminates a complex and changing relationship between irritability and psychopathology over development (Wakschlag et al., 2015b). This shift suggests several possibilities. First, this dynamic relationship may explain variability in specificity and effect size of the relation between irritability and psychopathology (Wakschlag et al., 2015b). Second, when irritability at preschool age was taken into account, decreased irritability added critical specificity to internalizing disorders at pre-adolescence. Finally, it is possible, although not directly examined here, that this transition to reduced irritability in school age may result increasing sophistication of self-regulatory capacities as well as the intersection of early irritability with environmental inputs (e.g., parenting, changing expectations of self-control in school environments) that interact with biological development (i.e., maturation of emotional brain regions), resulting in a compensatory drop in irritability behaviorally with the emotional tendency toward negative emotions remaining. Irritability emerges specifically as internalizing symptoms in pre-adolescence, rather than expressed irritability.

Developmental trajectories of irritability in school age predicted pre-adolescent gray matter volume a period when gray matter volumes peak (Giedd et al., 1999, 2012). Increased irritability in school age related to increased volumes of cognitive and social regions (e.g., cerebellum, temporal parietal junction/bank superior temporal sulcus, middle frontal gyrus, middle temporal, parietal regions), and decreased volumes in emotion/emotion regulation regions (e.g., amygdala, nucleus accumbens, orbitofrontal and ventrolateral frontal regions) in pre-adolescence after accounting for preschool age variability. This is consistent with a similar cross-sectional, longitudinal study of irritability (Pagliaccio et al., 2018), where early irritability changes (age 3-5 a combined PSA and ESA time point) related to cortical thickness four to five years later (7 to 10 years old). This study found that higher irritability related to increased thickness in left superior frontal gyrus, left superior temporal gyrus, and right inferior parietal lobule (Pagliaccio et al., 2018). That study, however, included a range of age cohorts, the irritability metric reflected a single additive dimension of irritability, and was oversampled for childhood symptoms of depression. In contrast, the current study examined the normal:abnormal spectrum of irritability longitudinally. The current findings are also consistent with extant studies that link adolescent internalizing and externalizing symptoms to variation in brain structure (Andre et al., 2019; Goodkind et al., 2015; Kaczkurkin, Park, et al., 2019; Pagliaccio et al., 2018; Snyder et al., 2017). This literature suggests that general psychopathology is related to variation in amygdala (Andre et al., 2019; Goodkind et al., 2015; Snyder et al., 2017), hippocampal (Andre et al., 2019; Goodkind et al., 2015; Snyder et al., 2017), temporal regions (Pagliaccio et al., 2018; Snyder et al., 2017), and ventral frontal regions (Pagliaccio et al., 2018; Snyder et al., 2017) which, in the current study, had the largest association with school age irritability.

This pattern of findings converges with a larger adolescent literature further supporting that alterations in gray matter volume may reflect transdiagnostic risk emerging in school age (Aoki et al., 2014; Cole et al., 2014; Davis et al., 2018; Haijma et al., 2013; Niendam et al., 2012; Shang et al., 2014; Wise et al., 2017). Furthermore, these patterned alterations in gray matter volume suggest that irritability during school age predicts neurodevelopmental outcomes at pre-adolescence (Kaczkurkin, Moore, et al., 2019). Reduced gray matter volume may result from early myelination of immature emotional structures or over-pruning, whereas increases in volume may suggest excessive arborization, under-pruning, or a delay myelination in attentional and cognitive areas (Giedd et al., 2012; Kaczkurkin, Moore, et al., 2019). It is also noteworthy that the relation between gray matter volume and school age accounted for variance related to preschool age irritability, that suggests that these findings may largely affect neuromaturation of gray matter rather than stable early features of brain organization.

While this study provides a demonstration for how developmental science methods and concepts compliment RDoC to provide a new and invaluable perspective, emerging work with this emphasis is nascent. Take home messages from the present investigation highlight three such points. First, research on emerging psychopathology needs to start early and occur often. Longitudinal studies beginning at birth and employing parallel brain and behavioral methods over time are crucial to pinpointing the contribution of developmental variation to psychopathology and its neural substrates (Wakschlag et al., 2019). These findings also emphasize a second point, centering on the importance of contextualizing clinical metrics in terms of their biological and behavioral developmental contexts (Mittal & Wakschlag, 2017). Future studies should further investigate the overlapping and distinct trajectories of irritability that interact in a critical, interactive periods of changing environments (e.g., school contexts and social expectations of behavioral control) and changing biology (e.g., maturing brains). Collectively these findings also suggest a third and final important consideration about developmentally appropriate and empirically informed treatments. There is a broad window for identification of risk for psychopathology, developmentally tailored targeted interventions; the current paper may suggest a more general intervention at preschool age and target specific risk for externalizing and internalizing symptoms based on changes in the irritability patterns.

This study has a number of strengths, but there are some relevant limitations. First, it is notable that the current study oversampled for irritability to examine emerging psychopathology, but it will be critical for future studies to replicate these patterns in population-based samples since this sample was clinically enriched. There is also a growing literature that suggests that irritability varies by age and context. As a result, there is some potential that the observed effects are the result of some unmeasured developmental trait or environmental context. For example, it is possible that early irritability is related to prolonged language delay in the preschool age time point that resolves at school age as language improves (Roberts et al., 2018), though it should be noted that less than 1% of low irritability individuals move to high irritability over a 9-month follow-up (Wakschlag et al., 2005, 2015a; Wiggins et al., 2020). Similarly, previous work in preschool age irritability has demonstrated that reductions in frontal, amygdala, and accumbens volumes can result from features of the environment, which may partially account for the volume findings (Demir-Lira et al., 2016). To assess broader environmental and developmental contexts future studies should include more time points and account for environmental context (e.g. stress, parenting) to elucidate how these developmental domains and contextual processes may influence changes in irritability and gray matter volumes. Although the current study selected specific developmental timepoints, there may be missed nuance in development due to these age choices, which emphasizes the need for large longitudinal studies to examine the open clinical question of how often and when measures are predictive of future psychopathology. Similarly, the current study relies on a single time point to examine gray matter volumes; future studies should examine multiple neuroimaging time points to examine how the trajectories of neurodevelopment may reflect concurrent trajectories of irritability. In particular, the present study importantly demonstrates *that* accounting for developmental patterning enhances specificity of clinical prediction, but it does *not* tell us whether this is true at specific developmental time points (e.g. transition from the rapid developmental change of early childhood). However, it should be noted that the current sample size is similar to or larger than samples studied in much of the extant literature on irritability (Andre et al., 2019; Bolhuis et al., 2019; Briggs-Gowan et al., 2019; Karim & Perlman, 2017).

## Acknowledgments

We would like to thank Hubert Adam and James Burns for their excellent analytic work and support on MAPS study. The MAPS Study was supported by NIMH awards to Wakschlag and Briggs-Gowan (1R01MH082830; 2U01MH082830; U01MH090301). Work on this paper was also supported by NIMH grant to Wakschlag (R01MH107652). VAM and KSFD work was supported by the National Institutes of Mental Health (Grant R01s. MH094650, MH112545–01, MH103231, MH112545, MH094650, R21/R33MH103231. We gratefully acknowledge contributions of Joel Voss, Ellen Leibenluft, Daniel Pine and James Blair to the neuroimaging portion of the study. We thank the MAPS families for their generous participation in this longitudinal study, enabling to joyfully “map” the development of the MAPS youth. We have no conflicts to disclose.

## Notes

### Competing Interest Statement

The authors have declared no competing interest.

